# Genome analysis of clinical pneumococcal isolates to decipher sequence types, antibiotic resistance mechanisms and virulence gene profiles

**DOI:** 10.1101/2020.07.21.211649

**Authors:** Sreeram Chandra Murthy Peela, Steffi Rose, Sujatha Sistla

## Abstract

**Introduction:** *Streptococcus pneumoniae* is a common cause of meningitis and pneumonia. Various studies have explored the relationship between genome and phenotypic characteristics like antibiotic resistance.

**Methods:** Four pneumococcal isolates from clinical specimens were included. DNA was extracted and genomes were sequenced in 2×101bp layout. Standard genome analysis tools and pipelines were employed to understand phenotypic characteristics.

**Results:** There was almost perfect agreement of genome-based predictions and phenotypic methods for antibiotic resistance. Novel sequence type (ST) was identified in one of the isolates. In total 29 virulence genes were common for all the isolates.

## Introduction

Globally, *Streptococcus pneumoniae* remains one of the commonest causes of meningitis and pneumonia. Despite the implementation of pneumococcal vaccine, it continues to affect young children and cause significant mortality in this susceptible population. One of the reasons preventing pneumococcal infections remain a major public health challenge is the ability of this pathogen to uptake and integrate foreign DNA into its genome via a process called natural transformation. Due to this, the genome of pneumococcus is highly dynamic and many international groups are engaged in understanding the genetic diversity of pneumococcus. While some global studies like Global Pneumococcal Sequencing (GPS) projects (1) use isolates from various geographic regions, studies involving single centers and a few isolates also continue as the costs of genome sequencing has been reduced considerably in the last few years.

To understand the diversity of pneumococci isolated from clinical specimens, our center planned to investigate genomes of a very few number of isolates as a pilot study. We attempted to optimise various techniques like DNA purification and analysis of raw sequences using automated pipelines.

## Materials and Methods

### Bacterial isolates

Four pneumococcal isolates from clinical specimens – two from sputum (REO1453 and REO7484) and one each from blood (P924) and ascitic fluid (SF621) collected during the study period were randomly selected. The serotypes and resistance profiles are shown in the table 1.

**Table 1:**
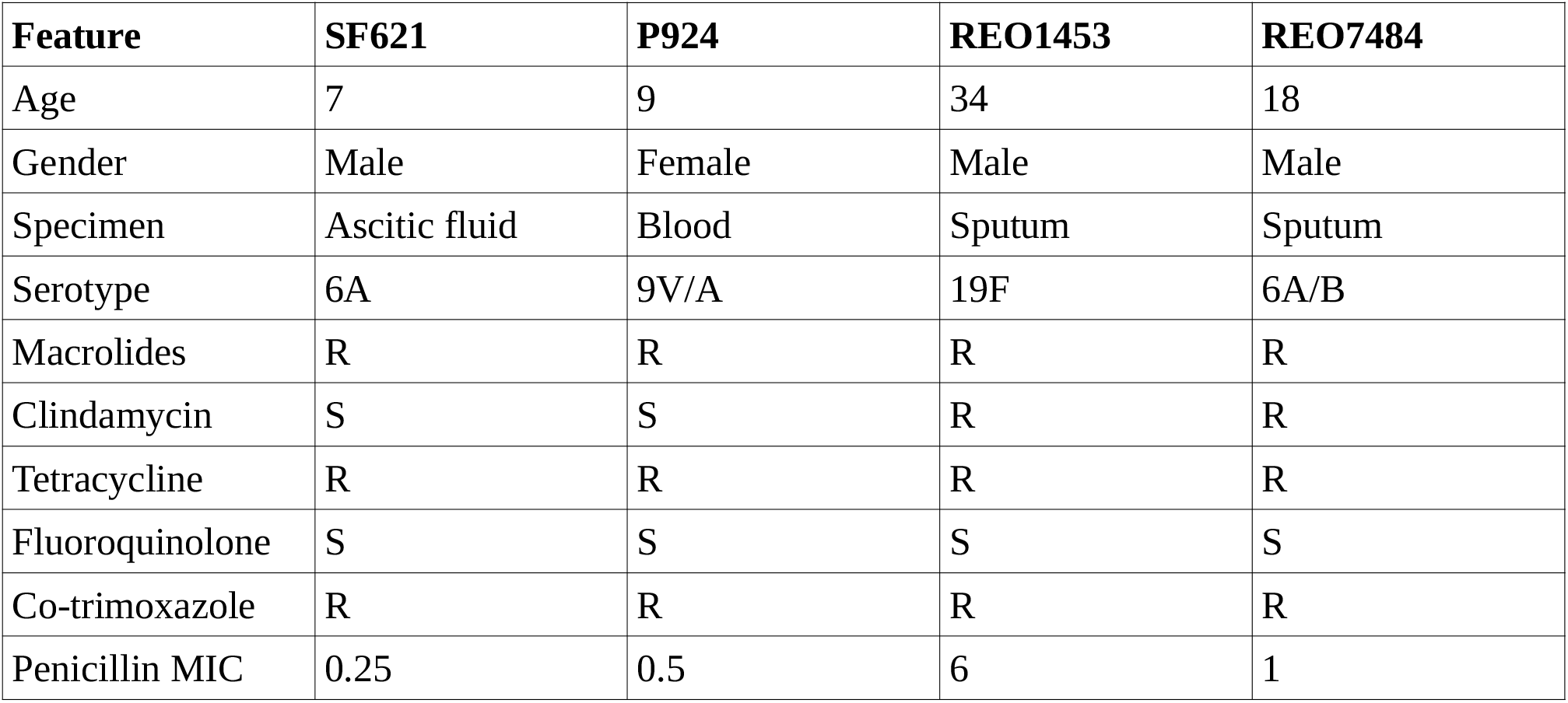
Features of the isolates selected for WGS. R – Resistant, S – Senstive. The penicillin MIC values are in μg/ml.

| Feature | SF621 | P924 | REO1453 | REO7484 |
| --- | --- | --- | --- | --- |
| Age | 7 | 9 | 34 | 18 |
| Gender | Male | Female | Male | Male |
| Specimen | Ascitic fluid | Blood | Sputum | Sputum |
| Serotype | 6A | 9V/A | 19F | 6A/B |
| Macrolides | R | R | R | R |
| Clindamycin | S | S | R | R |
| Tetracycline | R | R | R | R |
| Fluoroquinolone | S | S | S | S |
| Co-trimoxazole | R | R | R | R |
| Penicillin MIC | 0.25 | 0.5 | 6 | 1 |

### DNA purification

The selected isolates were cultured in BHI broth for 24 hours after which they were inoculated onto blood agar plates. The plates were further incubated for 24 hours in candle extinction jar at 37°C, and identified by using MALDI-TOF MS. DNA from these isolates was then extracted using QIAGEN Blood Mini DNA kit (Qiagen, Hilden, Germany) as per manufacturer instructions. The purified DNA samples were then checked for their quality using Nanodrop (the average of the values when repeated in triplicates was considered). Subsequently the samples were shipped to sequencing centre (Institute of Bioinformatics and Applied Biotechnology, Bengaluru, India) for whole genome sequencing.

### Library preparation and experimental set up

Quality of genomic DNA was analysed by Agilent Tape station 2200. About 200ng of DNA was taken and preceded for library preparation using NEBNext Ultra II DNA kit (E7645, New England Biolabs, Massachusetts, United States) as per the manufacturer’s instructions. The index sequences were mentioned in the table 2. Final Library was quantitated by Qubit 2.0 and fragment length was verified by Agilent Tape station 2200. Denatured libraries were clustered on cBot (Illumina) after which they were sequenced using HiSeq 2500 (Illumina) v3 chemistry. The expected read length was 101bp and in paired-end mode. Basecall files were demultiplexed to fastq formatted files using bcl2fastq v2.18.0.12.

**Table 2:**
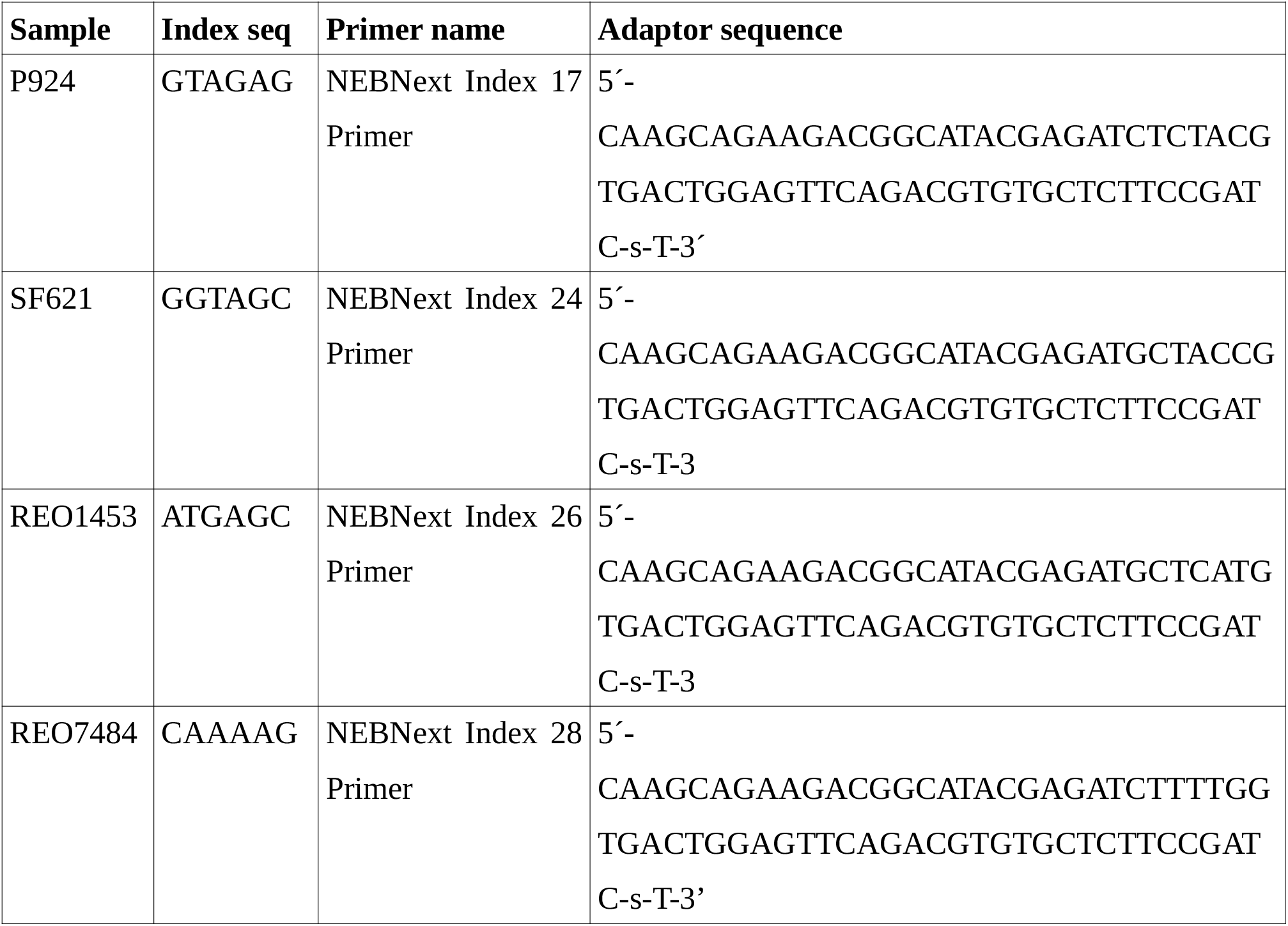
The primers, index sequences and the adapters used for the samples.

### Data analysis

The raw FASTQ files thus generated were assessed for quality using FASTQC v0.11.5 program (2). We also developed an automated shell script (available from https://github.com/sreerampeela/ngs_analysis/blob/assembly_annotation_V1/assembly_annotation.sh) for automating QC, assembly and annotation of genomes. Various parameters were selected based on best practices suggested by SPAdes Assembly Workflow using NextFlow pipeline (https://gitlab.com/cgps/ghru/pipelines/assembly) (table 3). Gene annotation and identification of virulence factors and antibiotic resistance genes were performed using Prokka and Abricate. The contigs were also annotated using Prokaryotic Genome Annotation Pipeline (PGAP) while submitting to NCBI GenBank. Simultaneously the SPN-TYPER program was used to predict the antimicrobial resistance, serotypes and MLST patterns from the FASTQ files (3–5). The MLST types were analysed using eBURST v3 (6). The predicted antibiotic resistance profiles and serotype patterns were then compared with the results obtained previously (table 3.8).

**Table 3:** Tools with version numbers and the parameters used for WGS analysis of the pneumococcal isolates.

| **Tool used** | **Parameters** | **Reference** |
| --- | --- | --- |
| Trimmomatic<br>0.36 | Min PHRED score:33, Illuminaclip: TruSeq2-PE.fa:2:30:10,<br>Leading:25, Trailing:25, Slidingwindow:4:15, Minlen:36 | (7) |
| MASH 2.1.1 | k-mer 32 and error k-mer 3 | (8) |
| SEQTK 1.3 | error max at 25 | (9) |
| LIGHTER<br>1.1.2 | -K 32 4920000 -maxcor 1 | (10) |
| FLASH 1.2.11 | min 20 max 100 | (11) |
| SPAdes 3.12.0 | K-mer sizes 21,33,43,53,63,75 | (12) |
| QUAST 5.0.2 |  | (13) |
| PROKKA<br>1.12 | GenBank files downloaded from NCBI | (14) |
| ABRICATE<br>0.5 | Argannot, CARD, ResFinder, NCBI (Antibiotic resistance genes)<br>VFDB (Virulence factors database) | (15) |

## Results and Discussion

All the four DNA samples had acceptable quality for WGS. The percentage of the integrated area was around 96.87 to 99.58 for the four samples. The raw reads obtained were submitted to NCBI SRA with Bioproject PRJNA532301 and the genomes were submitted to NCBI GenBank (accession ids in table 4 and the quality of assemblies were in table 5). The serotypes and antimicrobial resistance profile predicted the SPN TYPER program was in good agreement (except for a few changes in penicillin MIC predictions) with the results obtained in our laboratory (table 4). The new MLST pattern was submitted to PubMLST database and new sequence type 14608 was assigned. When the MLST patterns were analysed by eBURST algorithm, ST 14608 and 11921 were singletons while ST2697 belonged to group 2 and ST90 to group 1 (figure 1).

**Figure 1:**
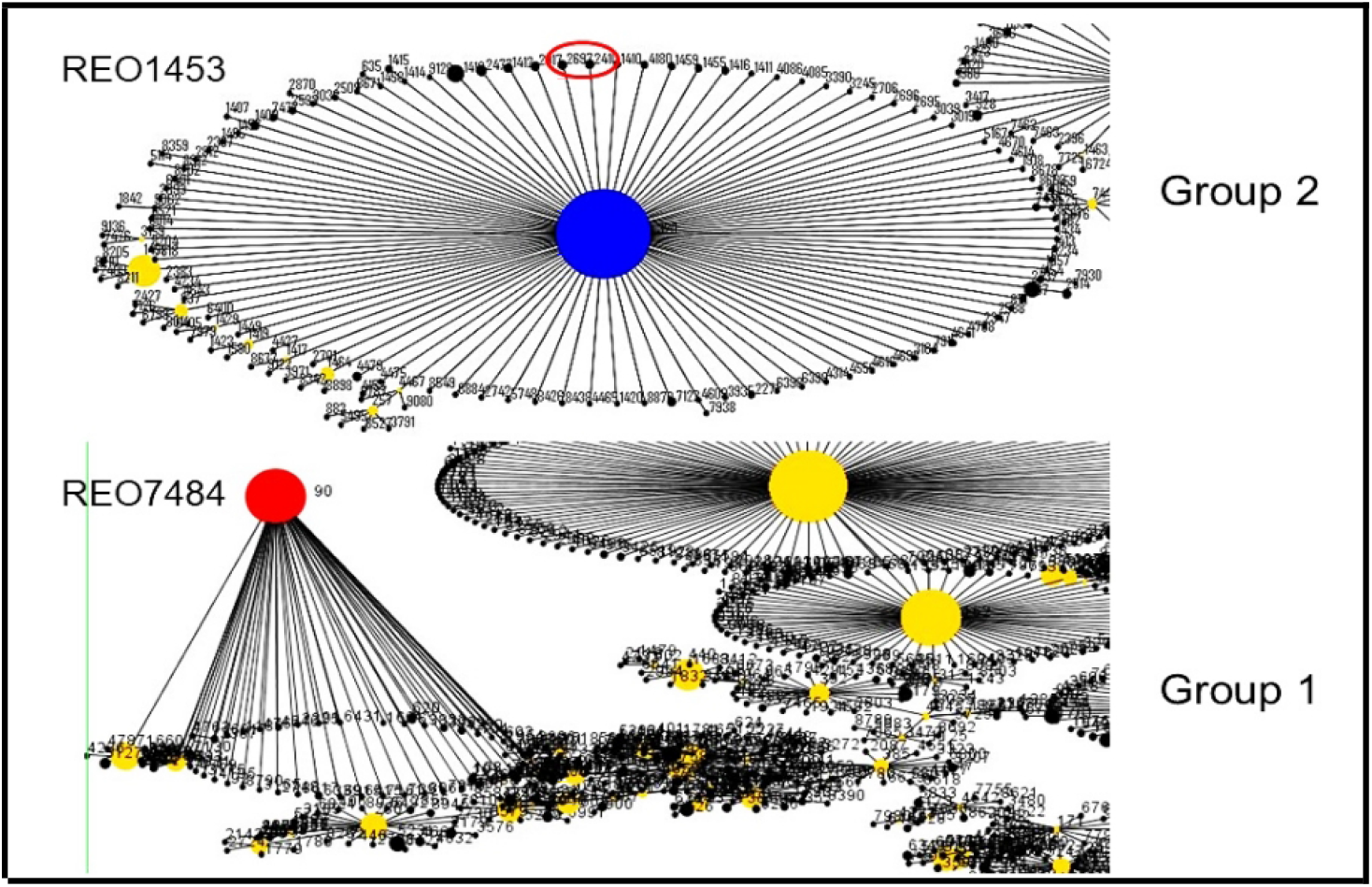
eBURST tree of the isolates REO1453 and REO7484.

**Table 4:** The results of SPN TYPER program. The AST profile was estimated based on the genes present. The new ST identified in the study is highlighted.

| **Feature** | **SF621** | **P924** | **REO1453** | **REO7484** |
| --- | --- | --- | --- | --- |
| Accession | VFBK000000000 | VFBI000000000 | VFBL000000000 | VFBJ000000000 |
| Serotype | 6A | 9V | 19F | 6B |
| Macrolides | R | R | R | R |
| Tetracycline | R | R | R | R |
| Fluoroquinolone | S | S | S | S |
| Cotrimoxazole | R | R | R | R |
| Penicillin MIC | 0.25 | 2 | 4 | 2 |
| aroe | 118 | 7 | 4 | 5 |
| gdh | 14 | 11 | 252 | 6 |
| gki | 44 | 10 | 16 | 1 |
| recP | 4 | 1 | 19 | 2 |
| spi | 1 | 6 | 15 | 6 |
| xpt | 6 | 667 | 6 | 3 |
| ddl | 20 | 1 | 20 | 4 |
| ST | **14608** | 11921 | 2697 | 90 |

**Table 5:** QUAST report for genome assemblies. All the samples had expected genome sizes and GC %.

| Statistic | SF621 | P924 | REO1453 | REO7484 |
| --- | --- | --- | --- | --- |
| # contigs ($\geq 0$ bp) | 309 | 498 | 469 | 613 |
| # contigs ($\geq 1000$ bp) | 73 | 82 | 69 | 76 |
| # contigs ($\geq 5000$ bp) | 53 | 54 | 52 | 63 |
| # contigs ($\geq 10000$ bp) | 47 | 43 | 43 | 51 |
| # contigs ($\geq 25000$ bp) | 31 | 29 | 24 | 29 |
| # contigs ($\geq 50000$ bp) | 15 | 19 | 17 | 15 |
| Total length ($\geq 0$ bp) | 2184789 | 2230068 | 2141461 | 2203480 |
| Total length ($\geq 1000$ bp) | 2122305 | 2117315 | 2044729 | 2115260 |
| Total length ($\geq 5000$ bp) | 2088062 | 2069048 | 2017057 | 2091944 |
| Total length ($\geq 10000$ bp) | 2045522 | 1993257 | 1959053 | 2017500 |
| Total length ($\geq 25000$ bp) | 1745359 | 1735782 | 1601474 | 1603272 |
| Total length ($\geq 50000$ bp) | 1160841 | 1395333 | 1331797 | 1062708 |
| # contigs | 95 | 119 | 101 | 89 |
| Largest contig | 163929 | 118980 | 167303 | 106329 |
| **Total length** | **2137828** | **2143173** | **2066110** | **2124423** |
| **GC (%)** | **39.78** | **39.87** | **39.88** | **39.49** |
| **N50** | **53718** | **57053** | **59285** | **53488** |
| N75 | 33599 | 31195 | 33834 | 28218 |
| L50 | 14 | 14 | 12 | 15 |
| L75 | 26 | 25 | 23 | 29 |
| # N's per 100 kbp | 0 | 0 | 0 | 0 |

The antimicrobial resistance genes and virulence factors were identified and annotated using ABRICATE with databases Argannot, CARD, ResFinder, NCBI and VFDB. Twelve antibiotic resistance genes were detected conferring resistance to penicillin, macrolides, tetracycline and other antimicrobials. Three isolates had similar number of antibiotic resistance genes while REO1453 had 10 antibiotic resistance genes. A total of 35 virulence genes were identified in the four isolates – 29 genes were common in all the four isolates while *cba* gene (encoding C-protein beta antigen) was detected in REO7484 and five genes in REO1453 (pitA-B, sipA, srtG1-2) (figure 2 and table 6). The PitA-B in REO1453 encode the minor and the major pili and were similar to a 19F serotype isolate from Taiwan. The two *srt*G1-2 genes encode a sortase while *sipA* is a signal peptidase. There were no major differences in the presence of the virulence genes among the isolates from different specimens.

**Figure 2:**
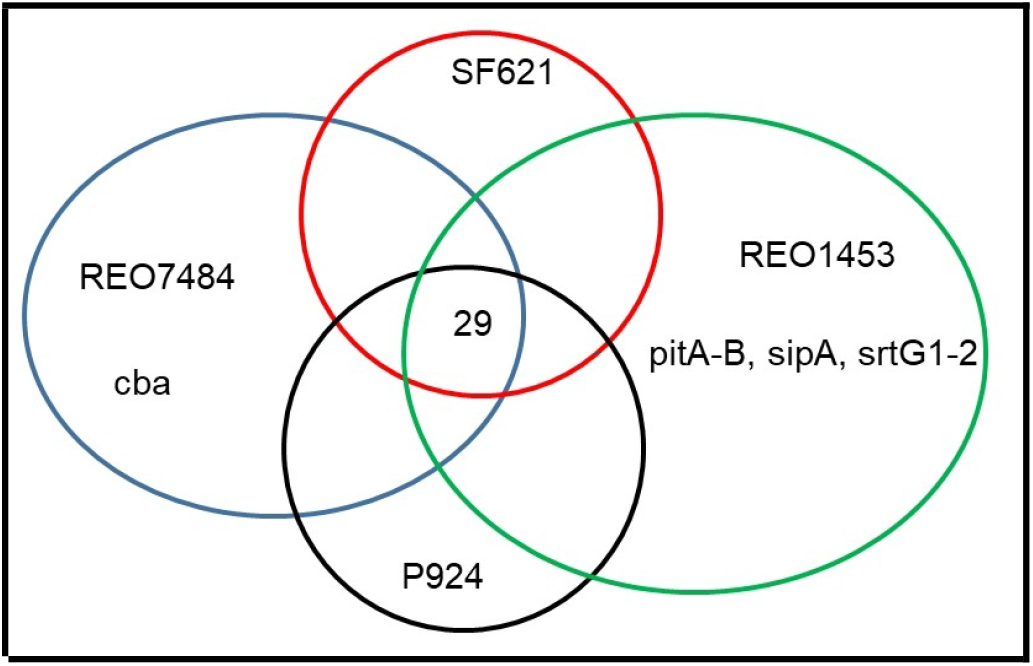
Distribution of virulence genes in the four tested isolates.

**Table 6:** The virulence genes detected among the four isolates.

| Gene | SF621 | P924 | REO1453 | REO7484 |
| --- | --- | --- | --- | --- |
| cba | N | N | N | P |
| cbpA | P | P | P | P |
| cbpD | P | P | P | P |
| cbpG | P | P | P | P |
| cps4A | P | P | P | P |
| cps4B | P | P | P | P |
| cps4C | P | P | P | P |
| cps4D | P | P | P | P |
| cpsB | P | P | P | P |
| hasC | P | P | P | P |
| hysA | P | P | P | P |
| lytA | P | P | P | P |
| lytB | P | P | P | P |
| lytC | P | P | P | P |
| nanA | P | P | P | P |
| nanB | P | P | P | P |
| pavA | P | P | P | P |
| pavB/pfbB | P | P | P | P |
| pce | P | P | P | P |
| pfbA | P | P | P | P |
| pitA | N | N | P | N |
| pitB | N | N | P | N |
| ply | P | P | P | P |
| psaA | P | P | P | P |
| pspA | P | P | P | P |
| pspC | P | P | P | P |
| rrgA | P | P | P | P |
| rrgB | P | P | P | P |
| rrgC | P | P | P | P |
| sipA | N | N | P | N |
| srtC-1/srtB | P | P | P | P |
| srtC-2/srtC | P | P | P | P |
| srtC-3/srtD | P | P | P | P |
| srtG1 | N | N | P | N |
| srtG2 | N | N | P | N |

### Insights and future directions

The objective of performing WGS for bacterial isolates is to obtain maximum genetic information with minimal costs. To understand the challenges in performing WGS for pneumococcal isolates, four samples were selected randomly in this study. We expected a significant difference in the patterns of virulence genes among the isolates as they were from different sources. The antimicrobial resistance patterns were almost similar and are the frequently identified combinations during the study period. To check for the MLST patterns, the WGS is now a viable alternative to normal PCR-based MLST deduction. Many of the tools used in this study are publicly available, and servers like Galaxy present all the tools at a single place to help in achieving speed and continuity during the analysis. Also, in the present study we developed and optimised a simple LINUX shell script to automate the genome assembly and annotation process and made it publicly available to facilitate people with minimal programming background to easily run the script. In future we plan to expand the genome analysis of pneumococci isolated from clinical specimens and derive valuable insights regarding pneumococcal genome dynamics.

## Funding

The study has been supported by JIPMER Intramural Research Fund and Senior Research Fellowship (SRF) from Council of Scientific and Industrial Research (CSIR), India.

## Acknowledgement

The authors would like to thank Dr. K.L. Ravikumar, Professor Emeritus and Chief - Central Research Laboratory (CRL), Kempegowda Institute of Medical Sciences (KIMS), Bengaluru for providing training in genome analysis.

## Notes

https://www.ncbi.nlm.nih.gov/bioproject/?term=PRJNA532301

### Competing Interest Statement

The authors have declared no competing interest.

